# Occurrence of Salmonella species in local and commercial chickens slaughtered at Samaru and Sabon-gari live bird markets, Kaduna State

**DOI:** 10.1101/2023.10.05.561012

**Authors:** Saaondo James Ashar, Anthony Oche Ameh, Jibril Adamu, Jacob Kwaga Paghi Kwaga

## Abstract

Salmonellosis is a bacterial infection caused by members of the genus *Salmonella*. It is one of the most common and important zoonotic diseases. The route of infection from animals to humans is usually through contaminated food, water and environment. Bacteriological investigation on 303 cloacal swab samples collected from local (139) and commercial (164) chickens slaughtered at Samaru and Sabon-gari live bird markets, Kaduna State was carried out to determine the occurrence of *Salmonella* species. Cultural isolation and identification using conventional biochemical tests, microgen test kit™ and Polymerase Chain Reaction (PCR) amplification of *inv*A gene of the *Salmonella* isolates was carried out. Isolates were subjected to antimicrobial susceptibility tests, phenotypic detection of Extended Spectrum Beta Lactamases (ESBLs) producing *Salmonella* isolates using the modified Clinical and Laboratory Standards Institute (CLSI) ESBL confirmatory test. The occurrence of *Salmonella* species based on cultural and biochemical tests was found to be 13 (4.29%), microgen kit identified 7 (53.8%) of the 13 *Salmonella* species while PCR based on *inv*A gene confirmed 9 (69.2%) as the overall occurrence rate of *Salmonella* species. Meanwhile, the study also revealed rate of occurrence of *Salmonella* species in local chickens 8 (61.5%) which was higher compared to commercial chickens 1 (7.69%) with a statistical significant association (**χ**^2^ = 8.775, P = 0.003) between the occurrence of *Salmonella* species in local and commercial chickens. Antimicrobial susceptibility testing was carried out using panel of 12 antibiotics from 8 different antibiotic classes. Highest rate of sensitivity of the isolates to antibiotics was observed for Ofloxacin with 90%, while all the isolates (100%) were resistant to Nalidixic Acid, Cefazolin, Amoxicillin + Clavulanic Acid, Oxacillin and Penicillin. This study revealed multidrug-resistance (MDR) profile of *Salmonella* with a total of (25%) of *Salmonella* isolates being resistant to 5 antibiotics belonging to 4 different classes of antimicrobials with the highest multiple antibiotic-resistant (MAR) indices ranging from 0.58 to 0.91. All the nine (9) isolates of *Salmonella* tested were found to be negative for ESBLs production. Therefore, it is concluded that *Salmonella* species are present in local and commercial chickens slaughtered at Samaru and Sabon-gari live bird markets, Kaduna State and could pose serious public health risks to handlers and to consumers of poultry meat and its products.

## CHAPTER ONE

### 1.0 INTRODUCTION

#### 1.1 Background of the Study

Foodborne diseases are caused by consuming contaminated foods or drinks. Myriad of microbes and toxic substances can contaminate foods. There are more than 250 known foodborne diseases (WHO, 2022). The majority are infectious and are caused by bacteria, viruses, and parasites (Adriana *et al*., 2017). Other foodborne diseases are essentially intoxications caused by toxins, or chemicals contaminating the food. All foodborne microbes and toxins enter the body through the gastrointestinal tract frequently causing nausea, vomiting, abdominal cramps and diarrhoea. The following are some of the examples of common foodborne diseases; campylobacteriosis (*Campylobacter* species), cryptosporidiosis (*Cryptosporidium* species), cyclosporiasis (*Cyclospora* species.), *Escherichia coli* O157:H7 infection (*Escherichia coli*), giardiasis (*Giardia* species), listeriosis (*Listeria monocytogenes*), salmonellosis (*Salmonella* species) (Maria *et al*., 2017).

The genus *Salmonella* is classified into two species, *Salmonella enterica* (type species) and *Salmonella bongori*, based on differences in their 16S rRNA sequence analysis (WHO, 2020). The type species, *S. enterica*, can be further classified into six subspecies based on their genomic relatedness and biochemical properties. The subspecies are denoted with Roman numerals: I, *S. enterica* subsp. *enterica*; II, *S. enterica* subsp. *salamae*; IIIa, *S. enterica* subsp. *arizonae*; IIIb, *S. enterica* subsp. *diarizonae*; IV, *S. enterica* subsp. *houtenae*; and VI, *S*. *enterica* subsp. *indica*. Among all the subspecies of *Salmonella*, *S*. *enterica* subsp. *enterica* (I) is found predominantly in mammals and contributes approximately 99% of *Salmonella* infections in humans and warm-blooded animals.

In contrast, the other five *Salmonella* subspecies and *S. bongori* are found mainly in the environment and also in cold-blooded animals, and hence are rare in humans (Alexandre *et al*., 2018).

Serotypes are groups within a single species of microorganisms, *Salmonella* bacteria look alike under the microscope but can be separated into many serotypes based on two structures on their surface: The outermost portion of the bacteria’s surface covering, called the O antigen; and a slender threadlike structure, called the H antigen, that is part of the flagella (WHO, 2021). *Salmonella* serotypes can be divided into two main groups: Typhoidal and Non-typhoidal. Non-typhoidal serotypes are zoonotic and can be transferred from animal-to-human and from human-to-human. They usually invade only the gastrointestinal tract and cause salmonellosis, the symptoms of which can be resolved without antibiotics (Alexandre *et al*., 2018). However, in sub-Saharan Africa, nontyphoidal *Salmonella* can be invasive and cause paratyphoid fever, which requires immediate treatment with antibiotics (Michael *et al*., 2018). Typhoidal serotypes can only be transferred from human-to-human, and can cause food-borne infection, typhoid fever, and paratyphoid fever. Typhoid fever is caused by *Salmonella* invading the bloodstream (the typhoidal form), or in addition spreading throughout the body, invading organs, and secreting endotoxins (the septic form). This can lead to life-threatening hypovolemic shock and septic shock, and requires intensive care including antibiotics (Jajere *et al*., 2019).

Infection with *Salmonella* is one of the most common and important zoonotic disease or infection that can spread between animals and humans. *Salmonella* can be transmitted both from animals to humans and *vice versa*. The route of infection from animals to humans is usually through contaminated food (NVRI, 2020).

When *Salmonella* species are ingested, they pass through a person’s stomach and colonize the small and large intestines. There, the bacteria invade the intestinal mucosae and proliferate. The bacteria can invade the lymphoid tissues of the gastrointestinal tract and spread to the bloodstream. Dissemination to the bloodstream depends on host factors and virulence of the *Salmonella* strain and occurs in less than 5% of infections. If the infection spreads to the bloodstream, any organ can become infected (e.g., liver, gall bladder, bones, or meninges) (CDC, 2019).

Multiple diseases can cause fever, diarrhoea, and abdominal cramps. Therefore, salmonellosis cannot be diagnosed on the basis of symptoms alone. To diagnose salmonellosis, the bacterium is usually isolated in the laboratory from the patient’s stool. The genus *Salmonella* is identified by using a series of biochemical tests. Subtyping (e.g., serotyping, pulsed field gel electrophoresis, and other tests) and antimicrobial susceptibility testing of *Salmonella* isolates are important adjuncts to the diagnostic testing of patients (Andrew *et al.,* 2017). Both provide insights into the epidemiology of the patient’s infection. Antimicrobial susceptibility testing also provides valuable information in the treatment of the patient, if use of antibiotics is deemed appropriate (Alexandre *et al*., 2018). The need for laboratory testing to diagnose infection with *Salmonella* affects our understanding of the occurrence of salmonellosis in the community. To be laboratory-confirmed: the patient must seek medical care, a specimen must be collected while the patient is still shedding the organism, and appropriate laboratory cultures must be performed (Ombelet *et al.,* 2019).

Infections due to *Salmonella* remain a global problem causing one of the most common foodborne illnesses (WHO, 2021). This condition in food animals play an important role in public health and particularly in food safety, as food products of animal origin such as eggs and eggs products, poultry meat, meat from other food animals and meat products are considered to be the major sources of human *Salmonella* infections (WHO, 2022).

In a bid to control salmonellosis, administration of antimicrobial agents in drinking water for treatment in poultry production has been widely used. However, research has shown an increasing resistance of *Salmonella* to antimicrobials frequently used in veterinary and public health practices (Economou and Gousia, 2015). Unfortunately, bacterial resistance to antibiotics which has been a reality almost since the dawn of the antibiotic era has within the past twenty years been compounded with an emergence of dangerous, resistant strains with a disturbing regularity (Jajere, 2019).

In addition, numerous countries have implemented National schemes to control *Salmonella* infections in animals in order to protect the consumer. The Zoonoses Directive 2003/99/EC in the European Union (EU) entails constant monitoring of breeding flocks of over 250 birds for *S.* Enteritidis and *S.* Typhimurium and destruction of positive flocks. Some other serotypes are also subject to special controls (Jajere *et al*., 2019).

Furthermore, administration of vaccines is becoming a more common means of control of *Salmonella* in poultry and importantly, immunity from such vaccinations should be differentiated from field infections for proper surveillance (Jajere *et al*., 2019).

#### 1.2 Statement of the Research Problem

Reports on the occurrence of *Salmonella* in local chickens in Nigeria is scarce to both live bird market workers and the public hence predisposing individuals to the risk of transmission of infection of salmonellosis (Bata *et al.,* 2019). Although, the incidence of *Salmonella* infections is on the increase worldwide, studies have not fully demonstrated comparisons between the occurrence of infection in local and commercial chickens in Nigeria. Hence, the type of bird at greater risk of infection is not known and this may be of public health implication to both live bird market workers and the entire populace at large (Olalekan *et al*., 2018).

The emergence of antimicrobial resistance in *Salmonella* strains is a serious health problem on the increase in Nigeria particularly in poultry due to indiscriminate and improper use of sub-therapeutic dose of antibiotics (Kabiru *et al.,* 2022). This poses a high risk of zoonotic disease with the transmission of Multi-Drug Resistance (MDR) *Salmonella* strains from poultry to both live bird market workers involve in the processing of poultry and the public who consume poultry products.

Globally, *Salmonella* is of economic importance. There are reports of approximately 1.4 million cases of *Salmonella* infections occurring annually with an economic burden of over $14 billion in the United States of America (WHO, 2021). The economic impact of salmonellosis in Nigeria is increasing every year as Nigeria is one of the developing countries with many economic demands from various sectors including health (Fagbamila *et al*., 2017). *It* is one of the leading causes of infection, and this has a direct impact on the Nigerian marketing of the respective food-producing animals and animal-derived food products. Poultry or chicken salmonellosis related to host adapted serovars remains a major constraint on poultry production in all parts of Nigeria such as Gallinarum and Pullorum. Farmers still experience great losses (due to mortality, morbidity and drop in egg production) caused by host adapted *Salmonella* serovars despite huge amounts spent on vaccination and medication (Fagbamila *et al*., 2017).

#### 1.3 Justification of the Study

The findings in this study will fill the knowledge gap on the current status of *Salmonella* organisms in local and commercial chickens in Kaduna State and in Nigeria. Hence, reducing the risk of zoonotic transmission of salmonellosis to both the live bird market workers and the entre populace at large.

Determining the susceptibilities of *Salmonella* isolates to antibiotics commonly used in human and veterinary medicine could help in better therapeutic administration to avoid multi-drug resistance in both veterinary and human health. Therefore, more efficient preventive measures could be employed in order to prevent and reduce salmonellosis especially to those more at risk of being infected (Schroeder, 2017).

This research is aimed at determining the occurrence of *Salmonella* species in local and commercial chickens slaughtered at Samaru and Sabon-gari live bird markets. The outcome will enlighten the live bird market workers and the public on the risk posed by infected birds and contaminated poultry meat and meat products

#### 1.4 Aim and Objectives

##### 1.4.1 Aim of the study

The aim of this study was to determine the occurrence of *Salmonella* species in local and commercial chickens slaughtered at Samaru and Sabon-gari live bird markets, Kaduna State, Nigeria.

##### 1.4.2 Objectives of the study

The objectives were:

i. To isolate and identify *Salmonella* species in local and commercial chickens slaughtered at Samaru and Sabon-gari live bird markets Kaduna State, Nigeria.
ii. To detect *inv*A gene in *Salmonella* isolates using PCR.
iii. To determine the susceptibilities of *Salmonella* isolates to antibiotics commonly used in human and veterinary medicine.
iv. To phenotypically detect Extended Spectrum Beta Lactamases (ESBLs) producing *Salmonella* isolates using the modified Clinical and Laboratory Standards Institute (CLSI) ESBL confirmatory test.

#### 1.5 Research Questions

i. Is *Salmonella* species found in local and commercial chickens slaughtered at Samaru and Sabon-gari live bird markets Kaduna State, Nigeria?
ii. Do the *Salmonella* isolates from chickens in the study area harbour *inv*A gene?
iii. Are the *Salmonella* species in local and commercial chickens susceptible to commonly used antibiotics in human and veterinary medicine?
iv. Are the *Salmonella* isolates producers of Extended Spectrum Beta Lactamases (ESBLs) based on the modified Clinical and Laboratory Standards Institute (CLSI) ESBL confirmatory test?

## CHAPTER TWO

### 2.0 LITERATURE REVIEW

#### 2.1 Salmonella

*Salmonella* is usually spread by the faecal-oral route. *Salmonella* is commonly found in the gastrointestinal tracts of animals, and has been commonly associated with foods such as raw meat, chickens, eggs, and dairy products (WHO, 2022). Salmonellosis is a disease caused by the bacteria *Salmonella*. It is usually characterized by acute onset of fever, abdominal pain, diarrhoea, nausea and sometimes vomiting. Chickens and other chicken products are believed to be the main vehicles for transmission of *Salmonella* (Antunes *et al*., 2016). The commercial chicken industry has a common turnover rate of forty-two days in a broiler setting which can lead to the contamination of sites such as litter, air, or feed. Chickens are also raised in close proximity with other birds, which increases the likelihood of horizontal transfer. *Salmonella* is recognized as one of the most common cause of foodborne infections worldwide, resulting in millions of infections and significant human death annually (WHO, 2022). *Salmonella* species are responsible for an estimated 93.8 million cases of foodborne diseases in humans and an average of 155,000 deaths annually worldwide (WHO, 2022).

#### 2.2 Taxonomy of *Salmonella*

Scientifically *Salmonella* classification is described under Domain: Bacteria, Phylum: Protobacteria, Class: Gammaprotobacteria, Order: Enterobacterales, Family: *Enterobacteriaceae,* Genus *Salmonella* and Species: *Salmonella enterica* and *Salmonella bongori* (John *et al*., 2015).

Six subspecies are differentiated within *S. enterica* based on their biochemical characteristics and genomic structure, a Roman numeral and a name are used for the designation of these six subspecies as follows: I, *S. enterica* subsp. *enterica*; II, *S. enterica* subsp. *salamae*; IIIa,*S. enterica* subsp. *arizonae*; IIIb, *S. enterica* subsp. *diarizonae*; IV, *S. enterica* subsp. *houtenae*, and VI, *S. enterica* subsp. *indica* V (Mutalib *et al*., 2015).

#### 2.3 The Genome of *Salmonella*

The generation of complete genome sequences provides a blueprint that facilitates the genetic characterization of pathogens and their hosts. The genome of *Salmonella enterica* serovar Typhi (*S.* Typhi) harbours ∼5 million base pairs encoding some 4000 genes, of which >200 are functionally inactive. Comparison of *S.* Typhi isolates from around the world indicates that they are highly related and that they emerged from a single point of origin ∼30,000-50,000 years ago. Evidence suggests that, as well as undergoing gene degradation, *S.* Typhi has also recently acquired genes, such as those encoding the Vi antigen, by horizontal transfer events (WHO, 2021).

The genomes of enteric bacteria are under intensive selective pressure because of factors that include competition within the normal flora, coping with fluctuating nutrient sources in the host and the environment, and pressure from the host immune system. Thus, we may expect the genomes of enteric bacteria to show signatures of these evolutionary pressures, and this appears to be the case. Scattered along the core genome are blocks of genes or, in some cases, single genes or gene remnants that have limited or no homology with the core genome. Furthermore, these genes can be unrelated or show significant divergence between different enteric species or even between members of the same species. These novel genes can, however, share some related features and common functions (WHO, 2021). For example, they might work together to enhance the virulence potential of a species. Such virulence-associated gene combinations are often referred to as “pathogenicity islands”. Examples include *Salmonella* pathogenicity islands 1 and 2 (SPI-1 and SPI-2, respectively), which contribute to invasiveness and the ability of *Salmonella* organisms to survive inside eukaryotic cells (Rafaela *et al*., 2017). These gene clusters often have GC content that differs from that of the core genome, suggesting that they have been acquired more recently and independently by a horizontal gene-transfer event. Indeed, these genes are frequently associated with genes from plasmids and phage and are sometimes integrated into redundant transfer RNA genes that can act as receptor sites for foreign DNA. Prophages or phage remnants are signature genes of the diverse, noncore regions of the enteric bacterial genome (Rafaela *et al*., 2017).

##### 2.3.1 The impact of the *Salmonella* genome in animals

*Salmonella* found in the environment, food, and animals. The two types of clinical manifestations were associated with *Salmonella* serovars, including invasive, life-threatening systemic disease, referred to as typhoid fever, and self-limited gastroenteritis caused by NTS found in foods, animals, and the environment. However, 5% of individuals infected with NTS develop bacteraemia and disease manifestations are substantially different between different serovars. NTS were found in asymptomatic food-producing livestock, including poultry, sheep, cattle, and swine, indicating that *Salmonella* persistence and carriage in livestock are very common, possibly since *E. coli* and *Salmonella* diverged from a common ancestor (Mauro *et al.,* 2022). Invasive non-typhoidal *Salmonella* (iNTS) disease, are not typically associated with diarrhoea but present as non-specific febrile illnesses with symptoms that are clinically indistinguishable from other febrile illnesses, and with higher case fatality than is seen with non-invasive infection. Multiple sets of *Salmonella* genes are involved in prolonged infection and persistence (John *et al*., 2015). Distinct sets of fimbriae contribute to the intestinal persistence and colonization in different animal species. Besides fimbrial adhesins, other adhesins, including the autotransporter adhesins MisL, SadA, and ShdA, the type I secretion system-secreted adhesins SiiE and BapA, and curli biogenesis (*csg*) adhesins, were found to play a role in colonization and persistence in mouse gastrointestinal tract. Recent works suggested that biofilm formation is involved in *Salmonella* gall bladder persistence (Mauro *et al.,* 2022). *Salmonella* gall bladder colonization triggers upregulation of the O-antigen capsule-encoding operon (*yihU-yshA* and *yihV-yihW*) in an *agfD*-independent manner, which is specifically required for biofilm formation on cholesterol gallstones. Iron is an essential nutrient for human and animals. A major host defense against infection is nutritional immunity, e.g., *via* sequestration of metals, including iron, to prevent pathogen growth (Gibson *et al*., 2022). The siderophore ABC transporter FepBDGC is responsible for primary ferric ion import in *Salmonella*. It was shown that the *fep* system together with the ferric-iron-binding siderophores enterobactin and salmochelin is required for persistent *Salmonella* infection in mice. Multiple nonsynonymous single nucleotide polymorphisms (SNPs) were found in global virulence regulators, including DksA, RpoS, HilD, MelR, and BarA, and metabolic pathways, providing an adaptative advantage during persistence in the host (Mauro *et al.,* 2022). Meanwhile, the mean genome-wide rate of nonsynonymous to synonymous substitutions (*dN*/*dS*) was less than 1 during the short-term evolution of *Salmonella*, indicating that the underlying substitution rate is subject to purifying selection. In contrast to the relatively stable core genome, considerable variation in composition of mobile genetic elements, including prophages and plasmids, was identified within the same clone in the course of an epidemic (Mauro *et al.,* 2022). All these changes contributed to clinically relevant differences in phenotype and virulence, further emphasizing the critical importance of integrated genotypic data sets in understanding of biological variability in *Salmonella* epidemiology (Gibson *et al*., 2022)

##### 2.3.2 Plasmids

Plasmids of *Salmonella enterica* vary in size from 2 to more than 200 kb. The best described group of plasmids are the virulence plasmids (50-100 kb in size) present in serovars Enteritidis, Typhimurium, Dublin, Cholerae-suis, Gallinarum, Pullorum and Abortus-ovis. They all encode spvRABCD genes involved in intra-macrophage survival of *Salmonella*. Most isolates of *Salmonella enterica* serovar Typhimurium contain a 90-kb virulence plasmid. This plasmid is reported to be mobilizable but non-conjugative. However, it has been determined that the virulence plasmid of strains LT2, 14028, and SR-11 is indeed self-transmissible. An increasing number of *Salmonella* infections are resistant to antibiotics, and many of the genes responsible for those resistances are carried by plasmids. Plasmids are important mediators of horizontal gene exchange, which could potentially increase the spread of antibiotic resistance (AR) genes (CDC, 2019).

##### 2.3.3 *Salmonella* genomic evolution in the environment

Salmonellae possess multiple traits that permit survival in a diverse set of environments, such as soils, sediments, waters, and plant surfaces. In order to survive in these environments, *Salmonella* must be able to overcome several stresses, including extremes of temperature, pH, salt/osmotic pressure, moisture, exposure to UV, and predators, to name a few (Mauro *et al.,* 2022). Long-term persistence of these pathogens in the environment has been documented. For example, *Salmonella* introduced into corn crop soil through naturally contaminated poultry litter was recovered one year later (Martinz *et al*., 2019). In surface waters from the Eastern Shore of Virginia, *Salmonella* with the same pulsed-field gel electrophoresis (PFGE) pattern was isolated over multiple years. Some mechanisms that *Salmonella* use for survival in the environment are similar to those used during an infection. Interestingly even some virulence factors have been shown to be important to environmental survival. As more naturally occurring isolates are recovered and sequenced from various environments, more unique adaptations for survival in those environments may be discovered (Huanli *et al.,* 2018).

#### 2.4 Features of *Salmonella*

*Salmonella* are straight rods usually motile with peritrichous flagella, facultative anaerobe, ferment glucose usually with production of gas (except *S.* Typhi and *S.* Dublin), do not ferment lactose, sucrose, salicin and urea, reduce nitrate to nitrite and most are phototropic. *Salmonella* multiply optimally at a temperature of 35 to 37°C, pH of 6.5-7.5 and water activity between 0.84-0.94. Salmonellae do not grow well at low temperatures. However, salmonellae are hardy and not always killed by freezing (WHO, 2022). Most salmonellae survive well in acidic foods (pH ≤ 4.6) and resist dehydration. *Salmonella* grows readily on blood agar, MacConkey agar, Bismuth Sulphite agar or Deoxycholate agar. They are chemo-organotrophic, obtaining their energy from oxidation and reduction reactions using organic sources (IOS, 2002).

Most *Salmonella* species produce H_2_S, which can readily be detected by growing them on media containing ferrous sulphate such as triple sugar iron (TSI) where the *Salmonella* is able to produce H_2_S from thiosulphate. The bacteria are sensitive to heat and will not survive at temperature above 70°C so it is sensitive to pasteurization, but resistant to drying even for years especially in dried faeces, dust and other materials such as feed and certain food (Moraes, 2016). The usual habitat of different *Salmonella* species and subspecies is the intestines of both cold- and warm-blooded animals. The bacteria can also be found throughout the natural environment. Sources of the organism include water, soil, factory surfaces, kitchen surfaces, animal faeces, raw meats, raw poultry and raw sea foods. *Salmonella* can survive for number of weeks in water and for years in soil if there are favourable conditions such as temperature, pH and humidity (Moraes, 2016).

#### 2.5 Salmonellosis

Salmonellosis is a disease caused by members of the genus *Salmonella* (WHO, 2022). It is an important cause of diarrhoeal illness in humans, responsible for approximately 1.4 million illnesses and 600 deaths annually in the United States (WHO, 2021). Much of what is known about the epidemiology of salmonellosis came from outbreak investigations. These investigations have determined that most human infections result from the ingestion of foods of animal origin that are contaminated with *Salmonella* species (WHO, 2021). *Salmonella* of animal origin cause an intestinal infection, characterized by sudden onset of fever, myalgia, cephalagia, abdominal cramps, nausea and vomiting. There is increased risk of exposure among those who work in abattoirs, poultry processing plants and those in contact with animals and their products (Mahendra *et al.,* 2021).

##### 2.5.1 Infections due to *Salmonella*

The ability of *Salmonella* species to cause human infection involves attachment and colonization of intestinal columnar epithelial cells and specialized microfold cell overlying the Peyer’s patches (Shu-Kee *et al*., 2015). Symptoms of salmonellosis include diarrhoea, abdominal pain, nausea, and vomiting lasting 1 to 7 days. The illness is generally self-limiting in healthy adults with a mortality rate of *<*1% (Andrews *et al*., 2015). In severe cases, infection may progress to septicaemia and death, unless the person is promptly treated with the appropriate antimicrobials. Presently fluoroquinolones, macrolides, and third-generation cephalosporins are the drugs of choice (Roberto *et al*., 2019). Individuals who are immuno-compromised, children, infants, and the elderly are most likely to require antimicrobial treatment. Infections with antimicrobial-resistant strains may compromise treatment outcomes thus resulting in increased morbidity and mortality. In rare instances, some individuals can develop chronic conditions including reactive arthritis, Reiter’s syndrome, and ankylosing spondylitis (Moraes, 2016). The infective dose for salmonellosis in adult humans is estimated to be in the range of 104 to 106 cells or higher, but can be as low as 101 to 102 cells in highly susceptible individuals. Symptoms observed may be different due to variation in the dose of inoculation, mechanisms of pathogenicity, virulence factors, age and immune response of the host (Ranjana *et al.,* 2019). Other than gastroenteritis, *Salmonella* may also cause extra intestinal infection like meningitis, osteomyelitis, pneumonia, colecystitis, peritonitis, pyelonephritis, endocarditis, pericarditis, vasculitis and chronic conditions like aseptic arthritis and Reiter’s syndrome (Ranjana *et al*., 2019)

#### 2.6 Epidemiology of *Salmonella*

The environmental persistence of *Salmonella* is a significant factor in the epidemiology of *Salmonella* in poultry by creating opportunities for horizontal transmission of infection within and between flocks*. Salmonella* Heidelberg can be isolated from contaminated litter after 7 months of holding at room temperature (Ventola, 2015). Huanli *et al*. (2018) reported that *Salmonella* Typhimurium can survive for 6 months in feed and 18 months in litter stored at 25°C. In addition, *Salmonella* can survive for 28 days on the surface of refrigerated vegetables (Huanli *et al*., 2018).

##### 2.6.1 Distribution of *salmonella* in poultry in Nigeria

*Salmonella* is widespread in the intestines of birds, reptiles and mammals. People can acquire the bacteria via a variety of different foods of animal origin. The illness it causes is called salmonellosis and typically symptoms includes fever, diarrhoea and abdominal cramps. Besides fever, infected patients may also develop myalgia, bradycardia, hepatomegaly, splenomegaly, and rose spots on their chest and abdomen. In endemic regions, approximately 15% of the infected patients develop gastrointestinal complications which include pancreatitis, hepatitis and cholecystitis. Haemorrhage is one of the most severe gastrointestinal complications, that occur as a result of perforation of Peyer’s patches, lymphatic nodules located at the terminal ileum, resulting in bloody diarrhoea. In addition, the ability of typhoidal *Salmonella* to survive and persist usually results in relapse in approximately 10% of the infected patients (WHO, 2018). In persons with poor underlying health or weakened immune systems, *Salmonella* can invade the bloodstream and cause life-threatening infections. Anyone can become infected with *Salmonella*. Groups at highest risk for severe illness include: children younger than 5 years, adults older than 65, people with weakened immune systems, such as people with HIV, diabetes, or undergoing cancer treatment. In some cases, diarrhoea may be so severe that the person needs to be hospitalized. In some cases, infection may spread from the intestines to the bloodstream, and then to other parts of the body. In these people, *Salmonella* can cause death unless the person is treated promptly with antibiotics (Michael *et al*., 2018).

In 2010, the incidence of fever caused by Non-Typhoidal *Salmonella* called enteric fever was estimated to be 22 million cases resulting in 200,000 deaths worldwide, predominantly in developing countries like Nigeria. Meanwhile, Nigeria was estimated to have a medium incidence (10–100 cases per 100,000 persons). The incidence and mortality rate of enteric fever vary from region to region, but the mortality rate can be as high as 7% inspite of antibiotic therapy. Invasive Non-Typhoidal *Salmonella* (NTS) is endemic in Nigeria with high occurrence rates in children below 3 years of age and human immunodeficiency virus (HIV)-infected patients, and these invasive strains confer a mortality rate up to 25%. The burden of foodborne diseases is substantial. Every day almost 1 in 10 people fall ill and about 20, 000 get infected every month. Foodborne diseases can be severe, especially for young children in Nigeria. Diarrhoeal diseases are the most common illnesses resulting from unsafe food. *Salmonella* is one of the four key global causes of diarrhoeal diseases (WHO, 2021) and these organisms have been reported as being the largest contributor to death statistics (39%) among all foodborne pathogens. As mentioned by CDC, the outbreak (2010) in USA involved the contamination of eggs by *Salmonella* Enteriditis, resulting in 1939 cases of NTS infections across 16 states (Abdurrahman *et al*., 2020).

The transmission of NTS infection to humans can occur through the ingestion of food or water contaminated with infected animals’ waste, direct contact with infected animals or consumption of meat and other products derived from infected food animals. The worldwide incidence rate of NTS infection is high as the strains can be found naturally in the environment and in both domestic and wild animals including cats, dogs, amphibians, reptiles and rodents (WHO, 2018).

In the early 1960s, the first incidence of *Salmonella* resistance to a single antibiotic, namely chloramphenicol, was reported. Since then, the frequency of isolation of *Salmonella* strains with resistance towards one or more antimicrobial agents has increased in many countries, including the USA, the UK and Saudi Arabia (WHO, 2018). Antimicrobial agents such as ampicillin, chloramphenicol and trimethoprim–sulfamethoxazole are used as the traditional first line treatments for *Salmonella* infections. *Salmonella* serotypes resistant to these three or more different classes of antibiotics are referred to as Multi-Drug Resistant (MDR). Multiple Antibiotic Resistant (MAR) index is calculated as the ratio between the number of antibiotics that an isolate is resistant to and the total number of antibiotics to which the organism tested. A MAR greater than 0.2 means that the high risk source of contamination is where antibiotics are frequently used (WHO, 2018).

A work carried out by Awanye *et al*. (2022) in Nigeria revealed that multidrug-resistant (MDR) profile of *Salmonella* with a total of 9/14 (64.3%) of *Salmonella* Enteritidis being resistant to 5 antibiotics which belongs to 3 different groups of antimicrobials with a multiple antibiotic-resistant (MAR) index of 0.21; while 3/11 (27.3%) of *Salmonella* Typhimurium were resistant to 11 antibiotics which belongs to 7 different groups of antimicrobials with a MAR index of 0.46. This phenomenon has raised concern among public health authorities regarding both clinical management and prevention of the infection (Michael *et al*., 2018). The emergence of *Salmonella* with antimicrobial resistance is mainly promoted by the use of antibiotics in animal feed to promote the growth of food animals, and in veterinary medicine to treat bacterial infections in those animals (Awanye *et al.,* 2022). Moreover, MDR *Salmonella* strains were found in some exotic pet animals such as tortoises and turtles, as well as their water environment, and this could result in a higher risk of zoonotic infections in humans through direct contact with these animals (Michael *et al*., 2018). Furthermore, in recent years, a shift in *Salmonella* serotypes in poultry and poultry production has been reported in different geographical areas, being particularly associated with the spread of well adapted clones (Autunes *et al*., 2016). The administration of antimicrobial agents in animals has been used for treatment which creates selection pressure that favours the survival of antibiotic resistant pathogens. Drug resistance has been shown to be the most important hazard of drug residues (Roberto *et al*., 2019). Resistant bacteria could cause disease that is difficult to treat and may also transfer the resistance genes to some other pathogens (Bata *et al*., 2016).

Research studies on the occurrence of *Salmonella* in local chickens in Nigeria is scarce hence creating a gap in the knowledge of the current status of the organism available to local farmers. Furthermore, there is a need for research in order to know whether local or commercial chickens at live bird markets are greater risk of salmonellosis and the public health implication to both live bird market workers and the entire populace at large (Olalekan *et al*., 2018).

In a cross -sectional study of 150 intensively reared and 150 backyard chickens conducted by Obi and Ike (2015) in Nsukka environs, 412 birds representing an occurrence of 4% were positive for *Salmonella* with 1.3% and 6.7% respectively in intensively reared and backyard reared birds from cloacal swab samples. The occurrence of *Salmonella* was significantly higher (P < 0.05) in backyard chickens than in intensively reared chickens. Similarly, Fashae *et al*. (2010) in a study to determine the prevalence and antibiotic resistance of *Salmonella* serovars from humans and chickens in Ibadan, Nigeria, in 2004-2007 recorded a prevalence of 11% from chicken faecal samples out of the 641 faecal samples that were collected from poultry farms in Ibadan (Fagbamila *et al*., 2017).

##### 2.6.2 Modes of transmission of *Salmonella*

The primary reservoir of salmonellae is the intestinal tract of humans and animals, particularly in poultry (WHO, 2018). The organisms are excreted in faeces from which they may be transmitted by insects and other creatures to various places such as to water, soil and kitchen surfaces. Eggs, poultry and raw meat products are the most important food vehicles of *Salmonella* infection in humans (Huanli, 2019). *Salmonella* infection is usually acquired through direct or indirect contact with colonized animals, as well as by the oral route, mainly by ingesting contaminated food or drink (Andrew *et al*., 2015). Any food product is a potential source of human infection. *Salmonella* can be transmitted directly from human to human or from animal to human without the presence of contaminated food or water, but this is not a common mode of transmission. The Centers for Disease Control and Prevention (CDC) estimates that 76 million people suffer foodborne illnesses each year in the United States, accounting for 325,000 hospitalizations and more than 5000 deaths (CDC, 2019). The WHO (2021) Foodborne Disease Burden Epidemiology Reference Group (FERG) reported an estimated 582 million cases of 22 different foodborne enteric diseases, among them *Salmonella* were the main enteric disease agent responsible for most deaths in 2010. Infections with *Salmonella* alone accounted for one billion dollars yearly, in direct and indirect medical costs (Wing *et al.,* 2019). Since *Salmonella* has several opportunities to be introduced to the host, there is an increased risk of *Salmonella* reaching the consumer. Salmonellae can be transmitted vertically to the progeny of infected breeder flocks and horizontally within and between flocks (Wing *et al.,* 2019).

Vertical transmission of paratyphoid salmonellae to the progeny of infected breeder flocks can result from the production of eggs contamination by salmonellae in the inside or on the surface. During oviposition eggshells are often contaminated with paratyphoid salmonellae by faecal contamination. The penetration of salmonellae into and through the shell and shell membranes can result in direct transmission of infection to the developing embryo or can lead to exposure of the chick to infectious salmonellae when the shell structure is disrupted during hatching (Busse, 1995).

Horizontal transmission can occur by direct bird to bird contact, ingestion of contaminated faeces or litter, contaminated water, farm handlers (through their clothing, hand, shoes), farm equipment, and a variety of other sources (Maria *et al.,* 2017). Yichao *et al*. (2018) reported that, horizontal transmission could occur when unexposed day-old chicks were raised together with infected day-old chicks. Similarly, ESFA, (2019) reported that rodents serve as a means of introducing *Salmonella* into poultry facilities. Birds could acquire *Salmonella* when they feed on the droppings of mice in the farm. Likewise, the faeces of mice or snakes can contaminate the water and the feeds of the birds, which could serve as a means of transmission to the birds; EFSA, (2019) in their study confirmed the role of mice as a means of dissemination of *Salmonella* when they isolated *Salmonella* serovars S. Concord, S. Kentucky, S. Mbandaka, S. Montevideo and S. Molade from mice caught from broiler poultry farm. Contaminated poultry house environments are identified as one of the major implications of paratyphoid Salmonellae. Since the chickens itself is a reservoir of *Salmonella,* control measures need to start from the hatchery (Busse, 1995).

#### 2.7 Risk Factors for the Transmission of *Salmonella* in Chicken

The physiology of *Salmonella* has contributed to the difficulty in controlling environmental contamination and transmission of the organism. Salmonellae are bacteria with several potential vehicles, vectors, and reservoirs within a chicken flock (Kirsten, *et al*., 2021).

##### 2.7.1 Feed as a Source of *Salmonella*

Poultry feed and feed ingredients are the most common source of *Salmonella* contamination and consequently serve as an indirect means of infection to people consuming poultry meat and poultry products. Feeds are contaminated either from feed mills or on farm during feed formulation, feeding or handling and subsequently spread to poultry mostly through ingestion. *Salmonella* have the ability to survive under prolonged periods in dry conditions like feeds and may be recycled in all production stages in commercial feed preparation (Parket *et al*., 2019). Several factors most especially ingredients used in preparing poultry feeds have been implicated to be the major source of contamination (Thirumalaisamay *et al.,* 2019).

In Nigeria, Obiageli, (2015) reported an incidence of 100% *Salmonella* from different feed samples in Abia State; this was attributed to hygiene practice in the farms; Chenz, (2017) also reported 15% isolation rate of *Salmonella* species from chickens feed in Calabar, Cross River State. He observed that the main route of contamination was the feed that attracted animals and insect vectors that are *Salmonella* carriers and serve as a vehicle for introducing *Salmonella* into chicken feeds (Miranda *et al.,* 2018).

##### 2.7.2 Poultry as a source of *Salmonella*

Among livestock production systems, *Salmonella is* more frequently isolated from poultry (chickens, turkeys, ducks, geese and pheasants) than from other animals (OIE, 2018). CDC, (2015) reported that public health problems associated with salmonellosis were of poultry origin.

Poultry and eggs are considered the most important reservoirs from which *Salmonella* is passed into the food chain and ultimately transmitted to humans (WHO, 2018). The levels of these pathogen in poultry can vary depending on country, production system and the specific control measures in place. Poultry can carry some *Salmonella* serovars without any outwards signs or symptoms of disease. *Salmonella* can contaminate eggs on the shell or internally, but the egg shells are much more frequently contaminated than the white or the yolk. Furthermore, egg surface contamination is associated with many different serotypes, while infection of the white or yolk is primarily associated with *S.* Enteritidis (WHO, 2018). Contaminated poultry and poultry-derived products, including meat and eggs are a major source of foodborne salmonellosis (Autunes *et al*., 2016).

*Salmonella* is able to remain viable in frozen products as well as foods stored at high temperatures for long periods, due to their marked ability to persist in a wide range of varying environmental conditions. *Salmonella* can enter the food chain at any point. Contamination can occur at several stages in the slaughter process of poultry like faeces during evisceration, surfaces on the production line or cross contamination from contaminated products. Particular contamination hot spots in the poultry slaughter process include scalding, evisceration and cutting, while chilling in a water bath enhances cross-contamination (Amelie *et al*., 2017). The principal site of multiplication of these bacteria is the digestive tract, particularly the caecum, which may result in widespread contamination due to bacterial excretion through faeces (Kabir *et al.,* 2010).

*Salmonella* can be introduced to a flock *via* environmental sources such as feed, water, soil, bedding, litter material, faecal matter, rodents or contact with other poultry. As *Salmonella* colonizes the gastrointestinal tract, the organisms are excreted in faeces from which they may be transmitted by insects and other animals to a large number of areas and are generally found in contaminated water with faeces. Humans and animals that consume polluted water shed the bacteria through faecal matter continuing the cycle of contamination. All the fractions of poultry production can be affected by *Salmonella* organisms, like hatchery, incubators, breeding facilities, commercial raising operations of layers and broilers, feed preparation units and factories, transportation systems commercialization of facilities and slaughter houses (WHO, 2018).

#### 2.8 Host adaptation

Primarily, when infecting its host, *Salmonella* exists in the intestinal tract as a gastrointestinal pathogen with limited duration and disease progression. However, some serovars have adapted to cause an invasive disseminated disease (Jorge, 2021). During this process, these serovars also have lost the ability to infect a broad range of hosts, becoming much more host adaptive (HA) or host restrictive (HR). Host-adaptive serovars tend to have one or two main animals that they naturally infect but are capable of infecting other hosts given the opportunity. Host-restricted serovars have one main host and rarely or never naturally infect a different host. Well-known examples of HR serovars include *Salmonella enterica* serovar Typhi, *Salmonella enterica* serovar Paratyphi A, *Salmonella enterica* serovar Gallinarum/Pullorum, and *Salmonella enterica* serovar Abortusovis. The disease caused by these serovars in their natural host is typically characterized by fever and septicaemia, with very little or no gastrointestinal disease. Similarly, HA serovars, such as *Salmonella enterica* serovar Choleraesuis and *Salmonella enterica* serovar Dublin, cause severe systemic disease in their natural host and humans which is also characterized by fever and septicaemia with little diarrhoeal symptoms. This is in contrast to most *Salmonella* serotypes, which exhibit a broad or unrestricted host range and cause severe gastroenteritis. In recent years, though, there has been an emergence of HA serovars in certain clones of some unrestricted-host-range serotypes which cause invasive disease mainly in immunocompromised patients (Jorge, 2021).

#### 2.9 Clinical Manifestations of Salmonellosis in Birds

The clinical manifestation of salmonellosis in birds are; poop droppings of a sulphur with yellowish green colour is very much a symptom of this microorganism, your bird may have a ‘fluffed up’ appearance which is the sign of an unwell companion, damages to vital organs such as liver, spleen, kidney or heart, dermatitis and signs of scratching more than usual, weight loss and diarrhea can occur. Arthritis is a symptom with pigeons, conjunctivitis in extreme cases, and excessive persistent thirst (Eric *et al.,* 2021).

#### 2.10 Pathogenesis of *Salmonella* species

Almost all strains of *Salmonella* are pathogenic as they have the ability to invade, replicate and survive in human host cells, resulting in potentially fatal disease. *Salmonella* displays a remarkable characteristic during its invasion of non-phagocytic human host cells (John *et al*., 2018), whereby, it actually induces its own phagocytosis in order to gain access to the host cell. When the bacteria enter the digestive tract *via* contaminated water or food, they tend to penetrate the epithelial cells lining the intestinal wall. *Salmonella* pathogenicity islands (SPIs) encode for type III secretion systems, multi-channel proteins that allow *Salmonella* to inject its effectors across the intestinal epithelial cell membrane into the cytoplasm (John *et al*., 2018).

Following the engulfment of *Salmonella* into the host cell, the bacterium is encased in a membrane compartment called a vacuole, which is composed of the host cell membrane. Under normal circumstances, the presence of the bacterial foreign body would activate the host cell immune response, resulting in the fusion of the lysosomes and the secretion of digesting enzymes to degrade the intracellular bacteria. However, *Salmonella* uses the type III secretion system to inject other effector proteins into the vacuole, causing the alteration of the compartment structure. The remodelled vacuole blocks the fusion of the lysosomes and this permits the intracellular survival and replication of the bacteria within the host cells (John *et al*., 2018). The capability of the bacteria to survive within macrophages allows them to be carried in the reticuloendothelial system (RES). *Salmonella* causes a wide range of human diseases such as enteric fever, gastroenteritis and bacteraemia (John *et al.,* 2018).

#### 2.11 Virulence genes of *Salmonella*

*Salmonella* pathogenicity is mediated by numerous genes such as *invA*, *spiC* and *pipD,* which code for effectors that induce successful host infection. Pathogenicity of *Salmonella* is expressed in three ways such as host cell invasion, intracellular survival and colonization (Katja *et al.,* 2022). Numerous virulence genes are essential for *Salmonella* pathogenesis and these genes are located on various elements of the genome including the chromosome, plasmids, integrated bacteriophage DNA, *Salmonella* pathogenicity islands (SPIs), and *Salmonella* genomic islands (SGIs). SPIs are large gene cassettes and only SPI-1 and SPI-2 (not all SPIs) encode a membrane-associated type III secretion system (T3SS) which secretes a pool of 44 effector proteins, that alter the functioning of eukaryotic cells in order to facilitate bacterial pathogenicity inside the cell. Previous studies reported that SPIs are acquired by horizontal transmission and vertically passed to new clones (Katja *et al.,* 2022). More than 20 SPIs have been characterized, with greater focus on SPI-1 and SPI-2 that function via encoded T3SS since they harbour host invasion and intracellular survival genes. Inside the host cell, SPI-2 expresses genes that are important in intracellular survival, proliferation, and persistence in internal organs such as the spleen and liver. *Salmonella* spp. use virulence genes and factors located in SPI-1 for cell invasion and to initiate its pathogenicity (WHO, 2021). The invasion A gene (*invA*) is one of the most studied virulence factors that is also used as a biomarker for *Salmonella* species (Cherrie *et al*., 2018). It contains sequences that are unique to the genus *Salmonella*. Invasion A is a factor in the outer membrane of *Salmonella* spp. that is responsible for entering the host epithelial cells in the intestines thus initiating infection. The *inv* locus in *S. enterica* serovar Typhimurium was characterized and it was reported that *invA* is essential in the display of virulence in the intestine (Thobeka *et al.,* 2019).

One of the most important genes is *iroB*, a Fur-regulated gene located in a large DNA region which is used in the detection of *S. enterica* subspecies *enterica*. Previous studies which detected typhoid and non-typhoid *Salmonella* by PCR used *invA* and *iroB* together with flagellar genes. Furthermore, *iroB* was used to detect *Salmonella* from blood in another study. The *iroB* gene is a member of the *iroA* (*iroBCDEN*) gene cluster which is responsible for the synthesis and transport of enterobactin, a siderophore produced by *Salmonella* spp. and is essential for iron uptake inside the host. Besides enabling bacterial iron uptake, expression of the *iroA* cluster also facilitates the host immune escape by interrupting macrophage homeostasis. The specific role of *iroB* is to encode glucosyltransferase which glucosylates enterobactin. Enterobactin glucosylation contributes to the virulence of the bacteria by preventing the host antimicrobial protein (lipocalin-2) from sequestering the siderophore (Autunes *et al.,* 2016).

#### 2.12 Diagnosis of salmonellosis

There are four steps for the recovery of *Salmonella* cells from a food matrix. First is the pre-enrichment, where buffered peptone water can be used. This is followed by enrichment in selective broth, such as Rappaport-Vassiliadis (RV) broth, Selenite Cysteine Broth(SC), or tetrathionate broth (TT). Thirdly the subsequent isolation is done on selective media such as xylose lysine deoxycholate (XLD), Brillant green agar, Bismuth sulphite agar or Hektoen agar (HA) and *Salmonella* Shigella agar. Finally, it is by nucleic acid recognition method or phenotypic molecular typing as reported (Okopi *et al.,* 2016).

#### 2.13 Public Health Significance of salmonellosis

With the increase in poultry meat and egg consumption, the dynamics of animal production and consumer exposure have changed leading to new challenges in limiting poultry borne zoonoses like salmonellosis. This has significant implication in Nigeria where poultry industry is among the fastest growing sector. Globalization, commercialization and distribution have made it possible for a contaminated foodstuff to affect the health of people in several countries at the same time. The identification of only one contaminated food ingredient may lead to the discarding of entire lot, causing economic losses to the production sector and restrictions for international trade. The loss in animal production and public health issues associated with salmonellosis has a substantial impact on the economy of several countries (WHO, 2018). *Salmonella* is mostly transmitted to humans, through contaminated food and water. In hospitals, person-to-person transmission may also occur. Among veterinarians and farm workers, transmission by contact with infected animals is possible. Cross contamination of poultry can occur in slaughter houses as well as during preparation of poultry products. *Salmonella* can also lead to severe conditions like sepsis and death especially in infants and immune-compromised adults (Amelie *et al*., 2017). Several cases of foodborne salmonellosis that originated from poultry products have been reported globally (Jakociune *et al*., 2014). The predominant serovars of *Salmonella*, having public health importance are mainly *S*. Enteritidis and *S*. Typhimurium (Amelie *et al*., 2017), but changes in the environment and poultry raising practices coupled with increased international trading of poultry and its products have led to changes in predominance of different *Salmonella* serotypes. Recent concerns in public health point of view is antibiotic resistant serotypes (Musa *et al*., 2019).

#### 2.14 Prevention and Control of salmonellosis in poultry

Salmonellosis is one of the most common foodborne bacterial infections worldwide. It is caused by host specific and non-host specific *Salmonella* in poultry. Host specific infections include Pullorum disease and Fowl typhoid, whereas non-specific include infection by *Salmonella* Enteritidis and *Salmonella* Typhimurium. *Salmonella* infection in poultry leads to huge economic loss. As salmonellosis can be transmitted to humans, the presence of *Salmonella* in poultry and poultry products also hinder international trade (CDC, 2019). The following are measures for prevention and control of salmonellosis:

i. Eggs should be cleaned and disinfected or fumigated before hatching.
ii. Day old chicks should be bought from *Salmonella*-free breeder flocks in which no evidence of *Salmonella* has been detected.
iii. Chlorinated drinking water should ideally be given to birds, if not provide water from a clean drinking source.
iv. Monitor the *Salmonella* status of poultry feed, because feed contaminated with *Salmonella* is known to be a source of infection for poultry. Heat treated feed with or without addition of other bactericidal or bacteriostatic treatments are recommended.
v. Vaccination: In some places, vaccines both live and attenuated have been used to control the infection. Vaccination can be used as an overall *Salmonella* control programme.
vi. Depending upon animal health, and risk assessment, culling is an option to manage infected breeder and layer flocks. Infected flocks should be destroyed or slaughtered and processed to minimize human exposure.\
vii. Litter of infected birds should not be reused. Used poultry litter, carcasses and other contaminated farm waste should be transported and disposed off in safe manner to prevent direct or indirect contact of humans, livestock and other poultry to *Salmonella*.
viii. Proper care should be taken in cleaning and disinfection of the poultry house and equipment.
ix. If disease is introduced, control should focus on eradication of disease through isolation and destruction of contaminated flocks, proper disposal of carcasses and disinfection of fomites.

#### 2.15 Bacteriophages-based Tool for *Salmonella* control

Bacteriophages have been used for controlling bacterial infections based on their specificity to the host bacteria. Bacteriophages kept stable in thermal conditions from 30 to 60 °C and pH ranges from 3 to 13 can suggest the possibility of using bacteriophages in variable conditions. Recently, bacteriophages as a biocontrol tool have gained great attention and are recognized as an alternative for antibiotics. Bacteriophage control technique has been applied for *Salmonella* in vivo and food samples. When lytic bacteriophages were applied to the chicken skin contaminated with *S. enterica* serovar Enteritidis, less than one log reduction was obtained at the MOI of 1 and no viable bacteria were observed at the MOI of 10^5^. A new virulent bacteriophage, F01-E2, was isolated for controlling *S.* Typhimurium. F01 belongs to *Myoviridae* with a double-stranded deoxyribonucleic acid dsDNA genome of 86.2 kb and a broad host range (Shuai *et al*., 2019).

## CHAPTER THREE

### 3.0 MATERIALS AND METHOD

#### 3.1 Study Area

The study was conducted in Samaru and Sabon-gari live bird markets, Kaduna State, North-West, Nigeria. The two markets are located in Sabon-gari Local Government Area of Kaduna State. The two live bird markets were recently improved by the Federal Government as part of the Avian Influenza Control Project (AICP) (Figure 3.1). The Local Government is located at latitude 11^0^N and longitude 7.7^0^E. The climate is tropical in Sabon-gari, which experience two seasons; dry season usually from November to March and the wet season from April through October (Aunnus *et al*., 2018).

**Figure 3.1:**
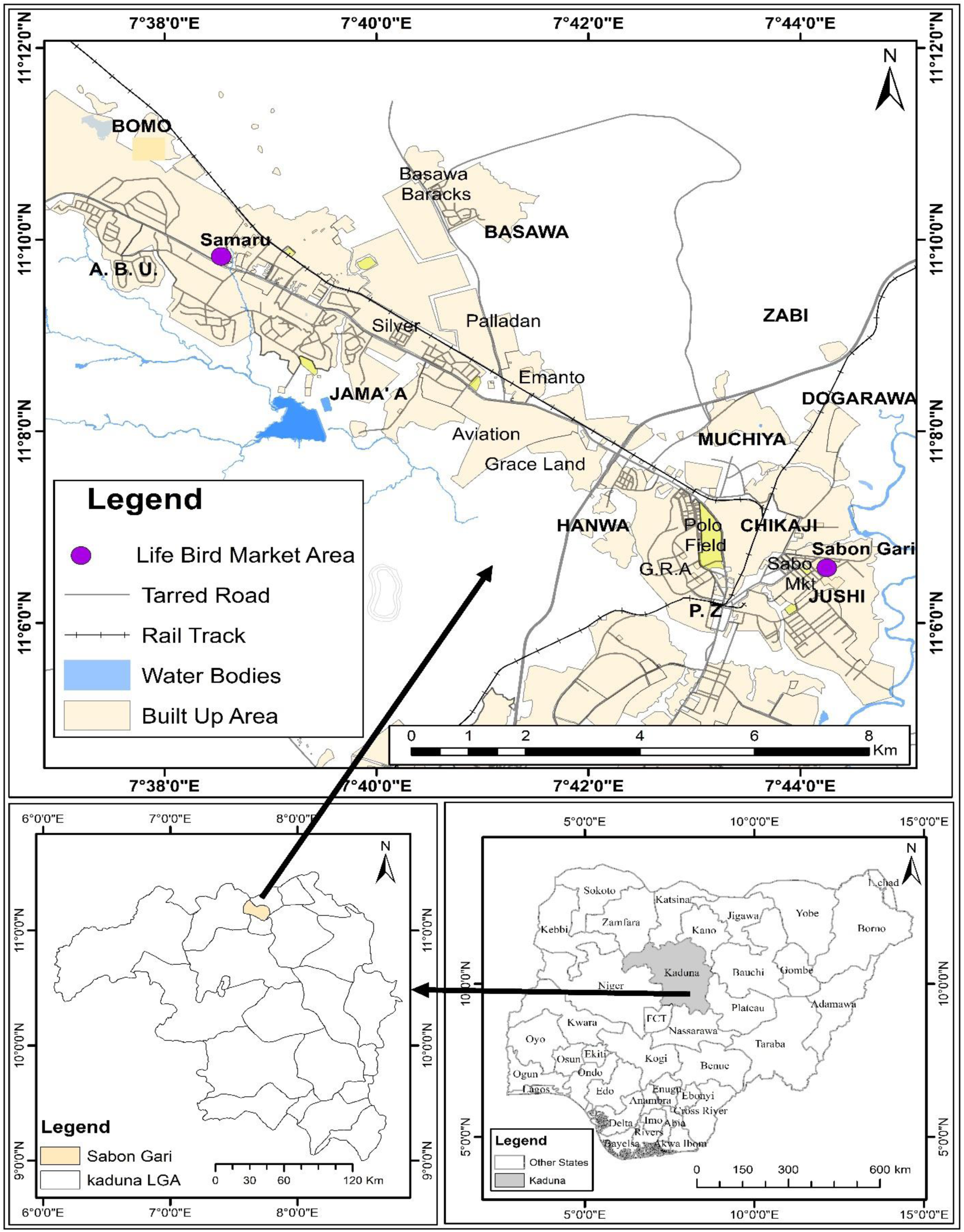
Map of Nigeria, Kaduna State, showing Sabon-gari Local Government Area and other Local Government Areas of the State. Source: <u>Open street map, 2023</u>

Poultry in the live bird markets (LBMs) are mainly from the small-scale commercial production system and village or backyard system, and occasionally from the commercial production system (Olubunmi *et al*., 2017). The LBMs are essential for sales of poultry in many developing countries, and they are a preferred place for many people to purchase poultry for consumption throughout the world (Olubunmi *et al*., 2017). Live Bird Markets worldwide serve as the most important mixing point of all birds, and for the maintenance and transmission of poultry diseases (Olubunmi *et al*., 2017). Several studies have linked human infection/outbreaks of influenza to LBMs (FAO, 2018). Biosecurity measures such as introduction of rest days, segregation of birds, cleaning and disinfection have been shown to significantly reduce the circulation of infection in the LBMs (FAO, 2018). Biosecurity requires the adoption of a set of attitudes and behaviours by people to reduce risk in all activities involving domestic, captive exotic, wild birds and their products (FAO, 2018). The state has vast expanse of fertile land growing both food and cash crops and it has rivers that provides opportunities for irrigation and fish farming. More than 70 percent of the work force earns their livelihood from the production of food crops, cash crops and livestock. Commercial poultry production is receiving wider popularity and acceptability by the day as a result of the growing demand for poultry meat and egg. Meanwhile, Kaduna State has an average total population of 8 million poultry production.

#### 3.2 Study Design

A cross-sectional study was used in determining the occurrence of *Salmonella* species in local and commercial chickens slaughtered at Samaru and Sabon-gari live bird markets, Kaduna State. The study lasted for 4 months and the sampling was purposive because they are two major markets in the study area.

#### 3.3 Sample Size Determination

The sample size was determined according to the formula by Thrusfield (2018):

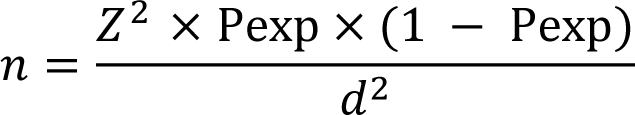

where:

n = Sample size

Z = appropriate value for the standard normal deviation for the desired confidence (1.96)

P_exp_ = previous prevalence

d = desired absolute precision (0.05)

- For local chickens, p= previous prevalence of 9% (Jasini *et al*., 2020)

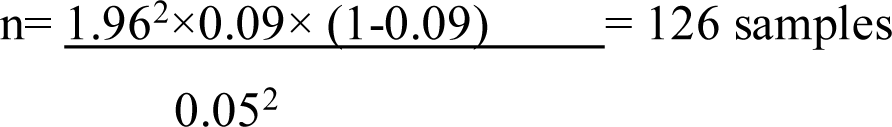
- For commercial Chickens, p=10.9% (Agada *et al*., 2014)

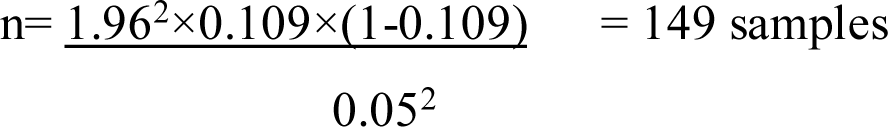
- Each of the calculated sample sizes were increased by 10% to give:
  ➢ Local chicken=139 samples
  ➢ Commercial chicken=164 samples

#### 3.4 Study Population

The study population is made up of local and commercial live chickens in Samaru and Sabon-gari areas of Kaduna State. The average daily slaughter of local chickens is 50/day in Samaru and 60/day in Sabon-gari. A total of 139 local chicken samples will be collected; 63 from Samaru and 76 from Sabon-gari. The average daily slaughter of commercial chickens is 200/day in Samaru and 250/day in Sabon-gari. A total of 164 commercial chicken samples were collected; 73 and 91 from Samaru and Sabon-gari respectively.

#### 3.5 Sample Collection and Transportation

Cloacal swab samples of local and commercial chickens were collected using sterile swab sticks at Samaru and Sabon-gari live bird markets, respectively for a period of 4 months from March to July, 2022. The normal saline moistened sterile swabs were inserted into the cloaca of the chickens and rotated round and was immediately inoculated into 3ml of Liquid Amies transport medium (Oxoid, CM111) for transportation to the Laboratory. The samples were maintained at 0 -4^0^C using ice packs in a cold box, transported to Department of Veterinary Public Health and Preventive Medicine and analyzed within 2-3 hours of collection at the Bacterial Zoonoses Laboratory of the Department.

#### 3.6 Isolation and Identification of *Salmonella* species from the samples

##### 3.6.1 Selective enrichment media

The 3ml suspension from Amies Transport medium was inoculated into 3mL of Rappaport Vassiliadis broth (Oxoid, CM0669) and was incubated at 37^0^C for 18-24 hours respectively (Busse, 1995).

**Table 3.1.** Distribution of slaughtered local and commercial chickens at the two sampling sites.

| Sampling site | Average daily slaughter |  | Calculated sample size |  |
| --- | --- | --- | --- | --- |
|  | Local | Commercial | Local | Commercial |
| Samaru | 50 | 200 | 63 | 73 |
| Sabon-gari | 60 | 250 | 76 | 91 |
| Total | 110 | 450 | 139 | 164 |

##### 3.6.2 Selective plating medium

Following enrichment, a loopful of culture from Rappaport Vassiliadis broth was sub-cultured by streaking onto Salmonella Shigella Agar (SSA) (TM Media, TM386) and incubated at 37^0^C for 18-24 hours (Busse, 1995). The plates were examined for the presence of typical colonies of *Salmonella* based on cultural and morphological characteristics by looking for smooth and opaque or colourless colonies which may have black spot in the centre due to hydrogen sulphide production on SSA.

##### 3.6.3 Conventional Biochemical Screening

Suspected colonies were screened for the following biochemical tests:

i. Triple Sugar Iron (TSI) test (Oxoid, CM 121): With a straight inoculation needle, a touch of the top of a well-isolated suspected colony of *Salmonella* was done. It was inoculated by first stabbing through the center of the medium to the bottom of the tube and then streaking the surface of the agar slant. It was capped loosely and slantly incubated at 37°C in ambient air for 18 to 24 hours. The typical *Salmonella* colonies produced alkaline slant over acid butt with or without gas or hydrogen sulphide gas.
ii. Urease test (Oxoid, CM 101): It was prepared by dissolving 2.95g of urea powder in 150ml of distilled water. It was then heated to dissolve and 5ml was dispensed into tubes and was allowed to cool and solidify in a slanted position. The suspected *Salmonella* colony was streaked, then capped and incubated at 37°C for 18-24 hour. It gave negative reaction as no colour change was observed.
iii. Simmons Citrate test (Oxoid, CM 110): It was also prepared by dissolving 2.86g into 100ml of distilled water. It was gently heated, with mixing, to boiling until agar was dissolved. It was then autoclaved and 5ml was dispensed into tubes slanted and allowed to cool and solidify. The suspected *Salmonella* colony was streaked aseptically, it was then capped and incubated at 37°C for 18-24 hour. Growth with color change from green to intense blue along the slant was observed as positive.
iv. Indole test (Oxoid, CM 120): It was prepared and dispensed and 5ml was dispensed into test tube. Suspected colony of *Salmonella* was streaked and incubated at 37°C for 18-24 hours. Then, a 0.5ml of Kavac’s reagent was added and absence of ring formation was observed for *Salmonella* isolates which gave a negative reaction.
v. Methyl red test (Oxoid, CM 132): Methyl Red and Voges Proskauer (MRVP) broth was prepared and 5ml was dispensed in test tubes. The suspected *Salmonella* colony was inoculated aseptically, it was then capped and incubated at 37°C for 18 -24 hours. A few drops of methyl red indicator were added in the incubated tubes and it gave positive reaction as the colour changed to a bright red
vi. Voges Proskauer test (Oxoid, CM 321): Methyl Red and Voges Proskauer (MRVP) broth was prepared and 5ml was dispensed in test tubes. The suspected *Salmonella* colony was inoculated aseptically, it was then capped and incubated at 37°C for 18-24 hours. A few drops of a solution A (alpha-naphthol) and solution B (Potassium hydroxide) were added. And it gave negative reaction for *Salmonella* as no pink red colour change was observed (Moraes, 2016).

##### 3.6.4 Identification of Presumptive *Salmonella* isolates by Microgen Bioproduct Limited (*GNA* MID-64CE) ^™^

Procedure for identification of *Salmonella* isolates using Microgen kit™

i. An emulsified colony from an 18-24 hours’ culture in 3ml normal sterile saline for the microwell test strip was prepared and mixed thoroughly.
ii. The adhesive tape was carefully peeled.
iii. Using a sterile Pasteur pipette, 2-3 drops (approximately 100μL) of the bacterial suspension was added to each well of the microwell test strip(s).
iv. After inoculation, the wells were overlaid and 2-3 drops of mineral oil were added in well 1,2 3 and 9. These wells were highlighted with a black circle around the well to assist in adding oil to the correct wells.
v. The top of the microwell test strip(s) were sealed with the adhesive tape removed earlier and incubated at 35-37°C.
vi. The microwell test strips were read after 18-24hours incubation for *Enterobacteriaceae*, for colour change as it was compared with a colour chart from the manufacturer of the kit (Surrey *et al.,* 2007).

#### 3.7 Extraction of DNA from *Salmonella* isolates

The bacteria were sub-cultured and stored on nutrient agar slant. DNA extraction was carried out by Phenol-Chloroform protocol (Barker,1998). A few colonies (2 or more) were picked and emulsified into 3ml normal sterile saline. A 200µLof the sample was added into 1.5ml tube and was mixed well; 400µL of lysis buffer was added to the sample contained in the 1.5ml tube and 10µl of proteinase K was added to the tube, and incubated at 55°C for 10minutes. A 400µL of equilibrated phenol was added, vortexed and centrifuged at 12000rpm for 5minutes. Supernatant was taken and 700µl of Chlorform (Isoamyl alcohol) was added, mixed well, and centrifuged at 12000rpm for 5minutes and it was transferred to a fresh tube. A 40µL of 3M sodium acetate was added. A 400µL of 100% ethanol was added, incubated at -20°C for 1hour, then, centrifuged for 15minutes at 4°C at 14000rpm to pellet the DNA. Supernatant was carefully removed without disturbing the DNA pellet. About 150µL of 70% cold ethanol was added, centrifuged at 4°C for 2 minutes at 14000rpm and the supernatant was discarded. The DNA pellet was dried at room temperature for 5 -10minutes and then the DNA pellet was resuspended in 100µL of TE buffer or molecular grade water by pipetting up and down. It was centrifuged briefly to collect the sample and the tube was placed on ice.

##### 3.7.1 PCR Amplification of *inv*A gene and reaction mixture

The quantification and purity of the extracted DNA was estimated using a Nano Drop spectrophotometer. The quantified DNA extract supernatant was prepared and used for PCR with *Salmonella* specific forward and reverse *inv*A primers S139 and S141, respectively, with the following nucleotide sequences based on the *inv*A gene of *Salmonella* forward primer 5’ GTG AAA TTA TCG CCA CGT TCG GGC AA-3’and reverse primer 5’ TCA TCG CAC CGT CAA AGG AAC C-3’ (Joseph *et al*., 2016). Reaction with these primers was carried out in a 20 μL amplification mixture consisting of 2.5 μL × 10x PCR buffer (500 mM KCl and 200 mM TrisHCl), 1.25 μL dNTPs (10 mM), 1.6 μL MgCl, 0.5 μL of each primer, 0.5 μL of Taq DNA polymerase (Fermentas), and 1.5 μL of DNA extract for each isolate was used in the reaction as template. Amplification was conducted in Master-gradient Thermo-cycler (Eppendorf).

##### 3.7.2 PCR cycling conditions

The cycling conditions were: an initial denaturation at 94°C for 60 s, followed by 35 cycles of denaturation at 94°C for 60 s, annealing at 64°C for 30 s, and elongation at 72°C for 30 s, and final extension for 7 min at 72°C. The PCR was carried out at Precision Biomedical Limited, Kaduna State.

##### 3.7.3 Electrophoresis

Agarose powder was measured (1g) and mixed with 100ml 1xTAE in a microwavable flask. It was microwaved for 1-3 minutes until the agarose was completely dissolved and it was allowed to cool to about 50°C. Then Midori green (4 μL) was added *to* bind to the DNA and allows visualizing the DNA under ultraviolet (UV) light. It was then poured into a gel tray with the well comb in place and it was maintained at about 4°C for 10minutes until it has completely solidified. It was placed into the electrophoresis unit after solidification. The gel box was filled with 1xTAE until the gel was covered. Molecular weight ladder was carefully loaded into the first lane of the gel and samples into the subsequent wells of the gel. Electrophoresis was carried out at 110V for 40 minutes and documentation carried out in a Gel documentation unit (Omega, HS90272000) using a UV light device for the visualization of the DNA fragments. The fragments of DNA are usually referred to as ‘bands’ due to their appearance on the gel (Begum *et al.,* 2010). The tank could only run eight (8) samples per row for electrophoresis, that is why the gel was divided into two. Lane 1 as positive control for the two gels.

#### 3.8 Antimicrobial susceptibility profiles of the isolates

The antimicrobial susceptibilities of *Salmonella* isolates were conducted using a modified Kirby-Bauer disk diffusion method (Bauer *et al*., 1966). A panel of 12 antibiotics commonly used in human and veterinary medicine belonging to 8 different classes of antibiotics were screened for Penicillin (Amoxicillin+ clavulanic acid 20µg + 10µg, Penicillin 10µg, Oxacillin 1µg), Chloramphenicol (Chloramphenicol 30µg), Aminoglycoside (Gentamicin 10µg), Cephalosporins (Cefazolin 30µg), Sulfonamide (Trimethoprim 5µg), Nitrofurans (Nitrofurantoin 30µg), Quinolones (Nalidixic acid 30µg), Fluoroquinolones (Ofloxacin 5µg, Enrofloxacin 5µg), Polymyxin (Colistin 10µg) were tested. The surface of each inoculum was touched with a sterile inoculating loop and transferred into 4ml of Mueller Hinton broth. The turbidity of the inoculum was adjusted using sterile saline solution to a 0.5 McFarland standard. The diluted bacterial suspension was streaked onto Mueller Hinton agar plates using sterile cotton swabs. The plates were seeded uniformly by rubbing the swabs against the entire agar surface and allowed to dry for about 10 minutes. Each antimicrobial impregnated disk was applied onto the surface of the inoculated plate by using a disk dispenser (Oxoid^TM^). All plates were incubated at 37^0^C for 18-24 hours. Growth inhibition zones and classification of isolates as susceptible, intermediate and resistant was done according to recommendations of Clinical and Laboratory Standards Institute (CLSI, 2020). A reference strain of *E. coli* ATCC 25922 was used as quality control. Multidrug resistance (MDR) was defined as resistance of the isolate to at least one antibiotics in three or more classes of antibiotics (Magiorakos *et al.,* 2012). Multiple antibiotic resistance (MAR) index for each of the isolates was calculated by the formula given by Krumperman, (1983).

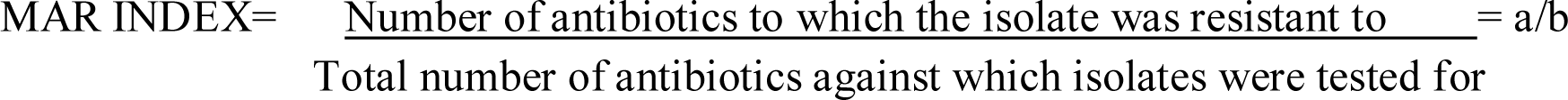

#### 3.9 Phenotypic detection of ESBLs of *Salmonella* isolates using the modified CLSI ESBL confirmatory test

A loopful of each isolate was pre-enriched in 5ml of buffered peptone water. A 0.3ml of ceftriaxone (30µg/ml) was added to a 200ml of Mueller-Hinton agar that was autoclaved and allowed to cool to about 55^0^C. Then, it was poured into 10 petri dishes for each of the 10 isolates and allowed to solidify. Sterile swab was dipped into the pre-enriched buffered peptone growth and streaked respectively onto the plates. A negative control of the Mueller-Hinton+ceftriaxone (30µg/ml) without the inoculum was used in order to check activity of the ceftriaxone. The plates were then incubated at 37^0^C for 18-24 hours. Growth of the isolates were observed on the test plates. The test was considered positive if growth was observed on the plates seeded with the isolates and negative for no growth (Salihu *et al*., 2020).

#### 3.10 Data Analysis

Statistical Package for Social Science (SPSS) version 20.0 was used for data analysis. Chi square was used to determine the association between the occurrence of *Salmonella* species and the type of chickens. Odds ratio and 95% confidence interval were used to test the strength of association between dichotomous variables. Occurrence rate was calculated using the formula:

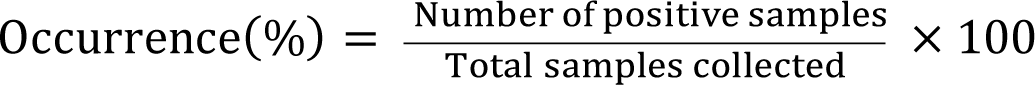

## CHAPTER FOUR

### 4.0 RESULTS

#### 4.1 Occurrence of *Salmonella* species in both local and commercial based on the cultural and Biochemical tests

The 303 cloacal samples collected from both local and commercial chickens were screened based on cultural and biochemical tests and 13 (4.29%) presumptive *Salmonella* species were identified. nine 9 (2.9%) were from local chickens while four 4 (1.3%) were from commercial chickens (Table 4.1).

#### 4.2 Phenotypic identification of *Salmonella* isolates using Microgen Biochemical Identification Kit (Bioproduct Limited, *GNA* MID-64CE) ^™^

The 13 *Salmonella* species based on cultural and biochemical tests were screened by Microgen Biochemical Identification kit and it detected seven as suspect for *Salmonella* species with inference ranging from 78.7% to 98% (Table 4.2).

#### 4.3 Polymerase chain reaction amplification of *inv*A gene in *Salmonella* species

The same 13 *Salmonella* species based on cultural and biochemical tests were tested for *inv*A gene by PCR, 9 (69.2%) were confirmed as *Salmonella* isolates (plates IA and IB). Chi-square was used to determine the association between occurrence of *Salmonella* species and type of chickens. Local chickens had a higher occurrence rate 8 (61.5%) compared to commercial chickens 1 (7.69%). There was statistical significant association (**χ**^2^ = 8.775, P = 0.003) between the occurrence of *Salmonella* species in local and commercial chickens (Table 4.3).

**Table 4.1:** Occurrence of *Salmonella* species in both local and commercial based on the cultural and Biochemical tests.

| **Type of Birds** | **No. of samples collected** | **No. of positive for *Salmonella* species** | **% of positive** |
| --- | --- | --- | --- |
| Local chickens | 139 | 9 | 2.97 |
| Commercial chickens | 164 | 4 | 1.32 |
| **Total** | **303** | **13** | **4.29** |

**Table 4.2:** Phenotypic identification of *Salmonella* isolates using Microgen Biochemical Identification Kit.

| S/N | Sample No. | Inference |
| --- | --- | --- |
| 1 | N16 | *Salmonella* spp. (78.7%) |
| 2 | N15 | *Salmonella* spp. (87.7%) |
| 3 | N17 | *Salmonella* spp. (98%) |
| 4 | N12 | *Salmonella* spp. (95.7%) |
| 5 | N11 | *Salmonella* spp. (92.3%) |
| 6 | C12 | *Salmonella* spp. (89.6%) |
| 7 | C14 | *Salmonella* spp. (97%) |

N = local chickens, C = commercial chickens)

**Plate IA:**
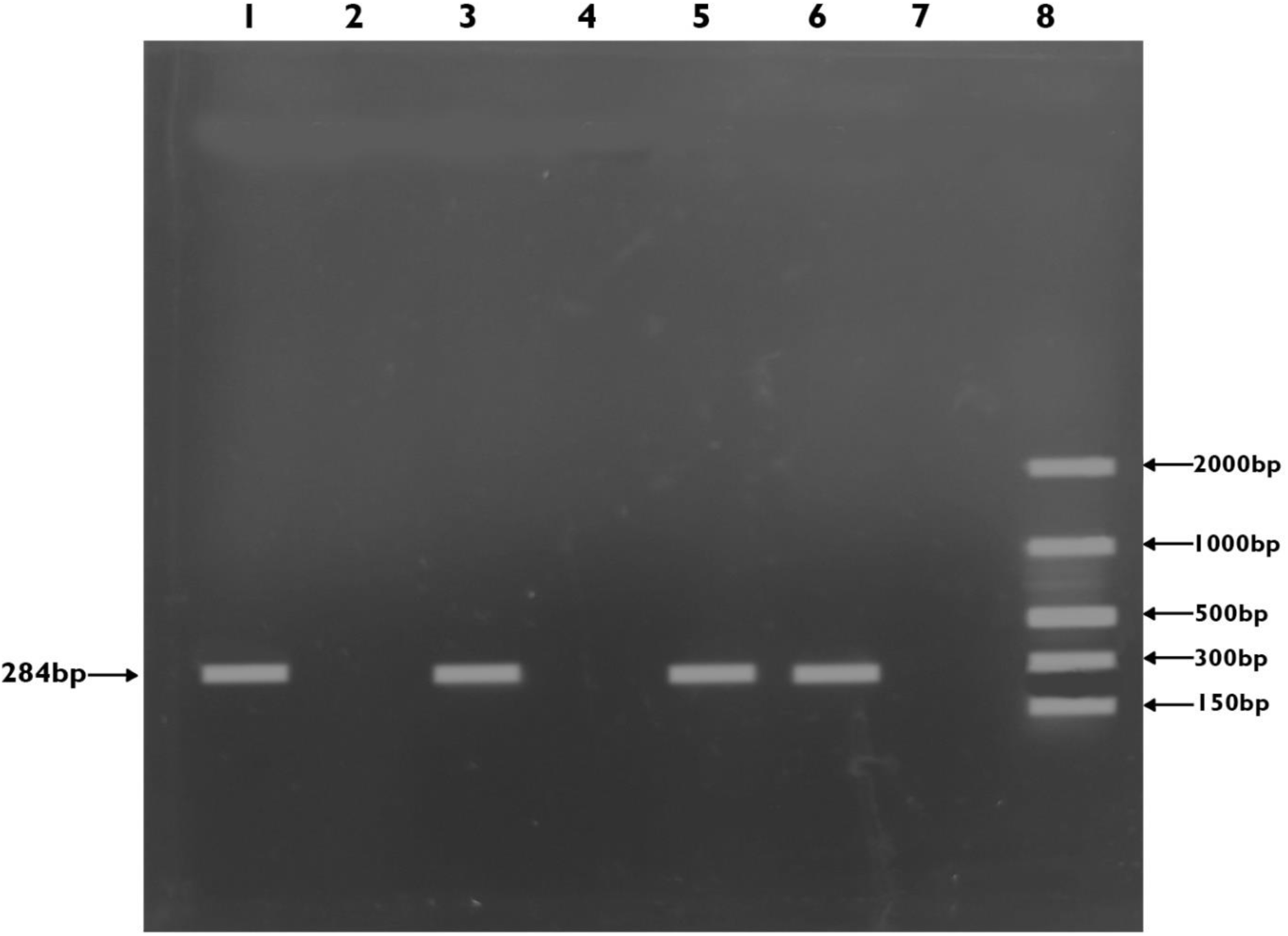
Gel documentation image of amplicons of *inv*A gene of *Salmonella* species. Lanes 3 (N4), 5 (N16), and 6 (N12) were positive for *inv*A while Lanes 2 (N1), and 4 (C17) showed no amplification (negative). Lane 1 was positive control, Lane 7 was negative control and Lane 8 Molecular weight marker, 100bp (BIONEER, 34302).

**Plate IB:**
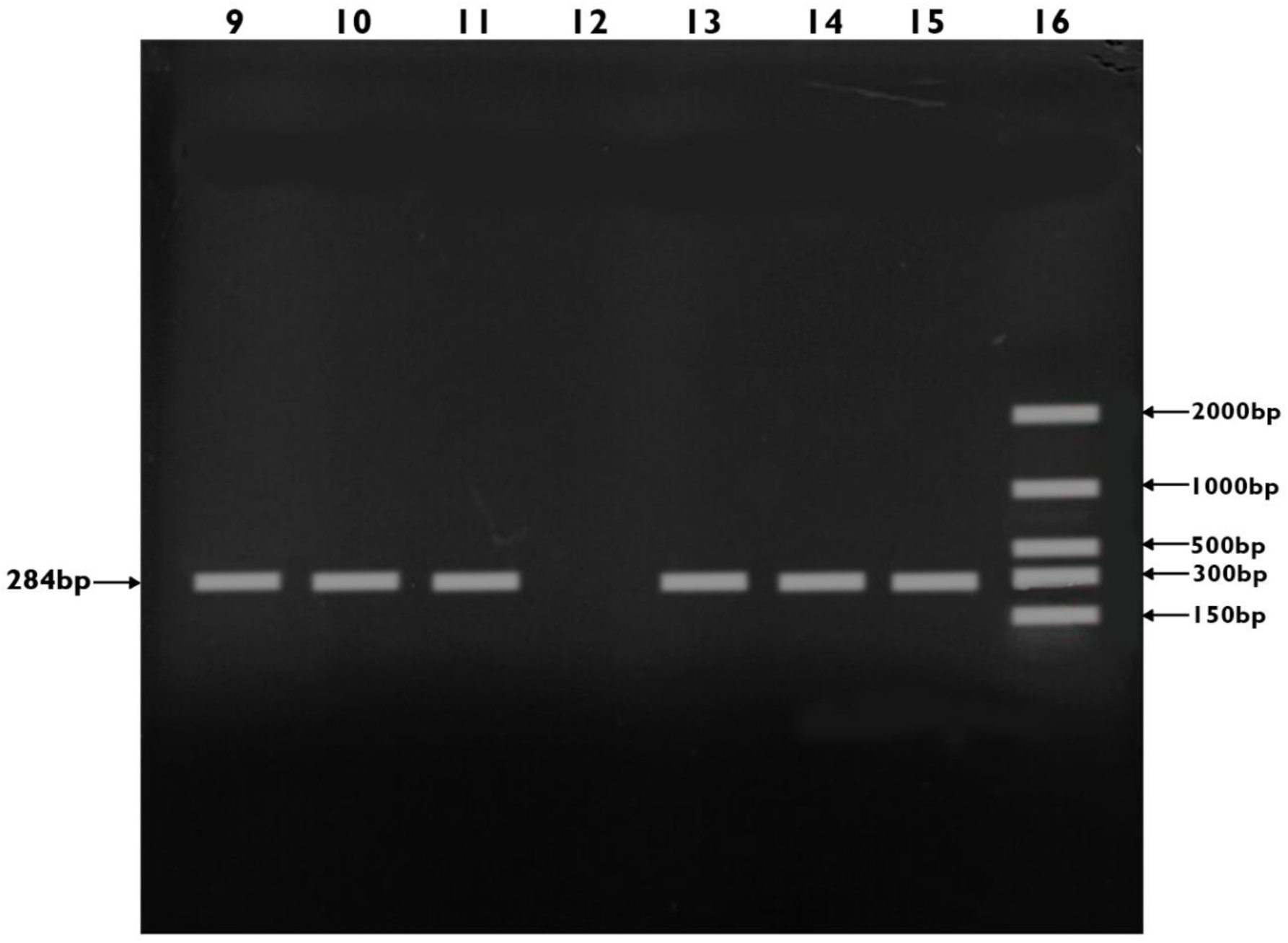
Gel documentation image of amplicons of *inv*A gene of *Salmonella* species Lanes 9 (N15), 10 (N11), 11 (N13), 13 (C14), 14 (N17) and 15 (N15) were positive for *inv*A while Lane 12 (C16) showed no amplification (negative). Lane 16 molecular weight marker, 100bp (BIONEER, 34302).

**Table 4.3:** Occurrence of *Salmonella* isolates in local and commercial chickens slaughtered at Samaru and Sabon-gari live bird markets, Kaduna State.

| **Type of Sample** | **No of samples collected** | **No. of positive for *Salmonella*** | **% of positive** | **Chi-Square ($\chi^2$)** | **P-value** | **Odds ratio(OR)** | **95% confidence interval on OR** |
| --- | --- | --- | --- | --- | --- | --- | --- |
| Local chickens | 9 | 8 | 61.5 | 8.775 | 0.003 | 0.1 | 0.016 – 0.642 |
| Commercial chickens | 4 | 1 | 7.69 |  |  |  |  |
| **Total** | **13** | **9** | **69.2** |  |  |  |  |

#### 4.4 Antimicrobial susceptibilities of nine (9) *Salmonella* isolates to 12 antibiotics

Antibiotic susceptibility testing was carried out on the nine isolates using the Kirby Bauer method. All the isolates (100%) were resistant to Cefazolin, Nalidixic Acid, Amoxicillin + Clavulanic Acid, Penicillin and Oxacillin (Table 4.4).

#### 4.5 Antibiogram profile of nine (9) *Salmonella* isolates

The antibiogram of *Salmonella* isolates is presented in Table 4.5. In this study, all 9 (100%) of the isolates were resistant to ≥3 antibiotics in different classes. In particular, isolate N12 was resistant to 11 out of 12 (91.6%) antibiotics. Similarly, isolates N04 and N11 were resistant each to 9 out of 12 (75%) antibiotics tested. Isolates C14, N16 and N15 were resistant each to 8 out of 12 (66.7%) antibiotics tested, while isolates C12, N02 and N17 were resistant each to 7 out of 12 (58.3%) antibiotics tested. The isolates had MAR index ranging from 0.58 to 0.91 being each resistant to 7-11 antibiotics. Meanwhile, there were four antibiogram (Table 4.5).

#### 4.6 Phenotypic detection of Extended Spectrum Beta Lactamases (ESBLs)

All the *Salmonella* isolates tested were found to be negative for extended spectrum beta lactamase production.

**Table 4.4:**
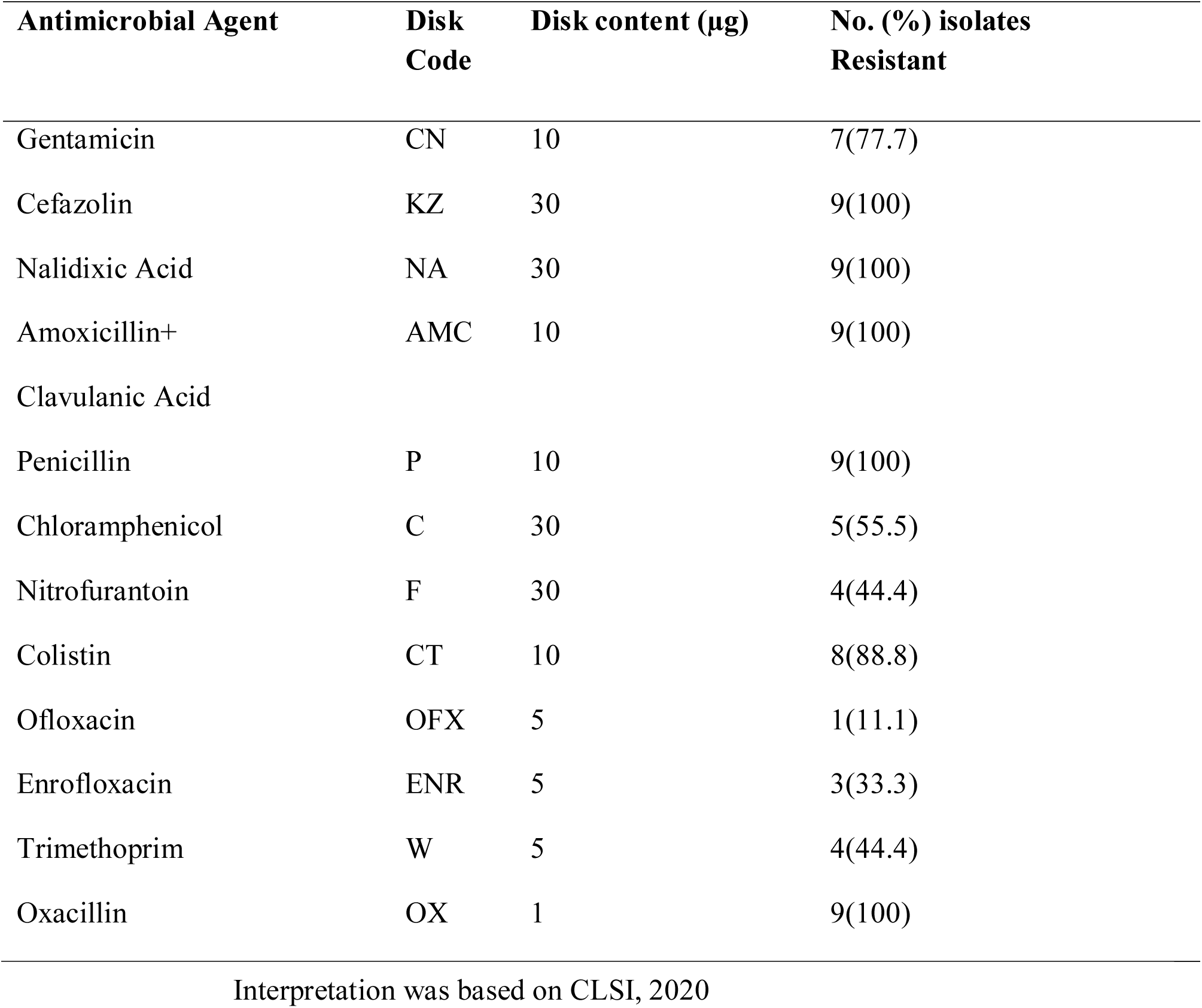
Antimicrobial susceptibilities of nine (9) *Salmonella* isolates to 12 antibiotics.

**Table 4.5:** Antibiogram of nine (9) *Salmonella* isolates tested against 12 antibiotics.

| S/N | Isolates No. | Resistance pattern of isolates tested | No. of resistant antibiotic agents | MAR index |
| --- | --- | --- | --- | --- |
| 1 | C12 | ENR, KZ, NA, AMC, P, OX, CT | 7 | 0.58 |
| 2 | N17 | CN, KZ, NA, AMC, P, OX, CT | 7 | 0.58 |
| 3 | N02 | CN, KZ, NA, AMC, P, OX CT | 7 | 0.58 |
| 4 | C14 | ENR, KZ, NA, AMC, P, C, OX, CT | 8 | 0.66 |
| 5 | N16 | CN, KZ, NA, AMC, P, F OX, CT | 8 | 0.66 |
| 6 | N15 | ENR, KZ, NA, AMC, P, C, OX, CT | 8 | 0.66 |
| 7 | N11 | CN, KZ, NA, AMC, P, C, F, OX, W | 9 | 0.75 |
| 8 | N04 | CN, OFX, KZ, NA, AMC, P, F, OX, CT | 9 | 0.75 |
| 9 | N12 | CN, ENR, KZ, NA, AMC, P, C, F, OX, CT, W | 11 | 0.91 |

Gentamicin (CN), Cefazolin (KZ), Nalidixic Acid (NA), Penicillin (P), Chloramphenicol (C), Nitrofurantoin (F), Colistin (CT), Ofloxacin (OFX), Enrofloxacin (ENR), Trimethoprim (W), Oxacillin (OX)

## CHAPTER FIVE

### 5.0 **DISCUSSION**

A total of 303 cloacal swab samples were collected from both local and commercial chickens at Samaru and Sabon-gari live bird markets. Based on cultural and biochemical tests, 13 (4.29%) *Salmonella* species were identified. This was lower than the occurrence rate of 10% reported by Jasini *et al*. (2020) but was similar to work documented by Awanye *et al*. (2022). This difference may have resulted from the differences in the production and handling of feeds, water, and biosecurity of the chickens in the various locations. The pathogenic *Salmonella* has been a major concern to public health all over the world. It is one of the major causes of bacterial food-borne gastroenteritis of humans associated with bacterial infection in poultry (WHO, 2021).

The 13 *Salmonella* isolates which were identified based on cultural and biochemical tests were subjected to both Microgen biochemical kit identification and PCR. Microgen kit identified 7 (53.8%) *Salmonella* isolates, while PCR based on *inv*A gene detection confirmed 9 (69.2%). This finding agrees with work documented by Mammam *et al*. (2014) with 58.4% by microgen and 70.2% by PCR and Fagbamila *et al*. (2017) with 48.6% by microgen and 62.4% by PCR who also confirmed higher rate of *Salmonella* isolates by PCR than Microgen kit in Kaduna and Plateau states respectively. This may be attributed to the fact that PCR is more specific and sensitive than microgen kit. Therefore, it is evident that *Salmonella* species are present in local and commercial chickens slaughtered at Samaru and Sabon-gari live bird markets, Kaduna State and could pose serious public health risks to handlers and to consumers of poultry meat and its products with possible zoonotic transmission of salmonellosis and resistant *Salmonella* strains from chickens to humans.

The overall occurrence rate of *Salmonella* species in both local and commercial chickens of 69.2% was higher than the occurrence of 31.7% reported by Musa *et al*. (2019) in Maiduguri. Meanwhile, there was higher occurrence of *Salmonella* species in local chickens compared to commercial chickens slaughtered at Samaru and Sabon-gari live bird markets, Kaduna State. Hence, there was association (χ^2^ = 8.775, P = 0.003) between the type of chickens slaughtered at Samaru and Sabon-gari live bird markets and *Salmonella* species infection. This is similar to findings by Ojo *et al*. (2012), who reported that free range indigenous chickens in Abeokuta, Nigeria had higher occurrence of *Salmonella* compared to commercial chickens, but contrary to the study by Jasini *et al*. (2020), in Maiduguri, Borno State. Thus, this may be as a result of the extensive system of management of local chickens, as they are allowed to roam freely and thereby exposing them to contaminated feed and drinking water. This poses a high risk of zoonotic disease with the transmission of *Salmonella* species infection from animals to humans *via* the ingestion of food or water contaminated with the animal faeces, direct contact or the consumption of the product from infected food animals. This finding indicates that *Salmonella* is associated with bacterial infection in chickens, the environment and that live bird markets can serve as potential hubs where disease agents are transmitted and maintained for prolonged periods of time.

The emergence of *Salmonella* isolates with antimicrobial resistance is mainly promoted by the use of antibiotics in animal feed to promote the growth of food animals, and in veterinary medicine to treat bacterial infections in those animals (Awanye *et al.,* 2022). Though, there is policy guiding the use of antimicrobials in animals in Nigeria, there is high level of indiscriminate use of antimicrobials in animals (Ojo *et al.,* 2012). The antibiotic susceptibilities of the isolates showed varying degrees of sensitivity. Greater proportions of the isolates were susceptible to ofloxacin 9 (90%), while all the isolates (100%) were resistant to nalidixic acid, cefazolin, amoxicillin + clavulanic acid, oxacillin, and penicillin. This finding is similar to the work of Fagbamila *et al*. (2017) in Plateau who reported that the highest level of resistance (100%) of the *Salmonella* isolates was to oxacillin, but contrary to that of Jasini *et al*. (2020) in Borno who reported 100% resistance of the *Salmonella* isolates to gentamicin. This may be as result of the choice and level of usage of the antibiotics in various locations. The indiscriminate and improper use of sub-therapeutic doses of antibiotics could also be responsible for the multidrug resistant isolates of *Salmonella* species (Eric *et al.,* 2021). This suggests that *Salmonella* species in local and commercial chickens at Samaru and Sabon-gari live bird markets, Kaduna State are multi-drug resistant and the chickens may serve as vehicles for transmission of resistant strains to humans. Therefore, the administration of these antibiotics in the treatment salmonellosis particularly to the resistant isolates in the study will not be effective. Meanwhile, Ofloxacin which is the most susceptible antibiotic to the isolates in the study could serve as choice antibiotic for treatment of salmonellosis at Samaru and Sabon-gari live bird markets, Kaduna State.

This study revealed antibiograms of all *Salmonella* isolates being resistant to a minimum of 7 antibiotics belonging to 4 different classes of antimicrobials giving multiple antibiotic-resistant (MAR) index ranging from 0.58 to 0.91. This agrees with work documented by Awanye *et al*. (2022) in Akure, whose findings showed that MAR index of *Salmonella* isolates were ≥0.2 but contrary to the work reported by Joseph *et al*. (2017) in Ogbomoso. This suggests that all the *Salmonella* isolates in this study may be from high risk of source of contamination either where antibiotics are frequently used or presence of resistant bacteria from the environment. The incidence of resistant *Salmonella* in Samaru and Sabon-gari live bird markets in Kaduna State, as observed in this study, may be due to indiscriminate usage of antimicrobials, the persistence and distribution of resistant bacteria, as well as the exposure of chickens to resistant bacteria in the environment. All these will influence the overall occurrence of antimicrobial resistant bacteria within the ecosystem (Ojo *et al.,* 2012). Although local chickens hardly receive any modern veterinary attention, they are exposed to potentially resistant bacteria harboured by other hosts (with previous exposure to antimicrobials) in the same environment (Ojo *et al*., 2012). Local chickens may acquire drug resistant *Salmonella* by contact with carriers or by ingestion of food and water that have been contaminated by faecal materials from other scavenging animals, that may have likely received veterinary care and treatment with antimicrobials. Poor sanitation and environmental pollution with human excreta due to inadequate toilet facilities, as observed in many rural communities in Nigeria, may expose local chickens which are free-rangers to resistant bacteria of human origin (Ejeh *et al*., 2017). Poor management of effluents generated from abattoirs and commercial farms also contribute to environmental pollution and hence, the possible exposure of free-range chickens to resistant bacteria. The multiple drug resistance of all the isolates raises serious public health concern because they could pose considerable health risk to both consumers and handlers of poultry meat product.

None of the nine (9) isolates of *Salmonella* species tested was found to be positive for ESBLs production. This is contrary to the work carried out by Rosangela *et al*. (2016) who detected extended spectrum beta lactamase from isolates in broiler processing plant in Brazil. This may be attributed to the methodology they used which is said to have 89.2% sensitivity and specificity of 100% for detecting ESBLs or it could be that the isolates have developed a different mechanism of expressing their resistance (Thobeka *et al.,* 2019).

## CHAPTER SIX

### 6.0 CONCLUSION AND RECOMMENDATIONS

#### 6.1 Conclusion

This study revealed 13 (4.29%) *Salmonella* isolates based on cultural and biochemical tests, Microgen kit detected 7 (53.8%) of the 13 *Salmonella* isolates while PCR based on *inv*A gene confirmed 9 as *Salmonella* isolates with overall occurrence rate of 69.2%

The study demonstrated that there was higher rate of occurrence of *Salmonella* infection in local chickens 8 (61.5%) compared to commercial chickens 1 (7.69%), with statistical association (χ^2^ = 4.850, P = 0.028) between *Salmonella* species infection and the type of chickens in live bird markets.

It was found that greater proportion of the isolates were susceptible to ofloxacin (90%) and all (100%) of the isolates were resistant to nalidixic acid, cefazolin, amoxicillin + clavulanic acid, oxacillin, and penicillin.

None of the nine (9) isolates of *Salmonella* species tested was found to be positive for ESBLs production.

#### 6.2 Recommendations

From the findings of the present study, the following recommendations are made:

i. Live bird market workers should ensure that they implement proper and personal hygiene in handling chickens in order to prevent zoonotic *Salmonella* infection.
ii. There should be public enlightment to sellers and buyers of birds on the possible occurrence of antibiotic resistant *Salmonella* in chickens.
iii. Measures should be taken to ensure that there is compliance with legislation on antibiotic use, in order to curtail incidence of Multi-Drug Resistance within the ecosystem of Samaru and Sabon-gari live bird markets of Kaduna State.
iv. Further studies should be undertaken to identify the serotypes and genotypic characterization of *Salmonella* isolates in local and commercial chickens slaughtered at Samaru and Sabon-gari live bird markets.

